# Opposite effects of nutrient enrichment and herbivory by an alien snail on growth of an invasive macrophyte and native macrophytes

**DOI:** 10.1101/2021.08.03.455002

**Authors:** Yimin Yan, Ayub M.O. Oduor, Feng Li, Yonghong Xie, Yanjie Liu

**Affiliations:** Key Laboratory of Wetland Ecology and Environment, Northeast Institute of Geography and Agroecology, Chinese Academy Sciences, Changchun 130102, China; University of Chinese Academy of Sciences, Beijing 100049, China; Dongting Lake Station for Wetland Ecosystem Research, Institute of Subtropical Agriculture, Chinese Academy of Sciences, Changsha, 410125, China; Department of Applied Biology, Technical University of Kenya, P. O. Box 52428 00200, Nairobi, Kenya; Ecology, Department of Biology, University of Konstanz, 78464 Konstanz, Germany

**Keywords:** Biological invasion, alien species, non-native plants, herbivory, shallow lake, invasion debt, early establishment, invasional-meltdown

## Abstract

Human-mediated introduction of plant and animal species into biogeographic ranges where they did not occur before has been so pervasive globally that many ecosystems are now co-invaded by multiple alien plant and animal species. Although empirical evidence of invaders modifying recipient ecosystems to the benefit of other aliens is accumulating, these interactions remain underexplored and underrepresented in heuristic models of invasion success. Many freshwater ecosystems are co-invaded by aquatic macrophytes and mollusks and at the same time experience nutrient enrichment from various sources. However, studies are lacking that test how nutrient enrichment and co-invasion by alien herbivores and plant species can interactively affect native plant communities in aquatic habitats. To test such effects, we performed a freshwater mesocosm experiment in which we grew a synthetic native macrophyte community of three species under two levels of nutrient enrichment (enrichment vs. no-enrichment) treatment and fully crossed with two levels of competition from an invasive macrophyte *Myriophyllum aquaticum* (competition vs. no-competition), and two levels of herbivory by an invasive snail *Pomacea canaliculata* (herbivory vs. no-herbivory) treatments. Results show that herbivory by the invasive snail enhanced above-ground biomass yield of the invasive macrophyte. Moreover, the invasive herbivore preferentially fed on biomass of the native macrophytes over that of the invasive macrophyte. However, nutrient enrichment reduced above-ground biomass yield of the invasive macrophyte. Our results suggest that eutrophication of aquatic habitats that are already invaded by *M. aquaticum* may slow down invasive spread of the invasive macrophyte. However, herbivory by the invasive snail *P. canaliculata* may enhance invasive spread of *M. aquaticum* in the same habitats. Broadly, our study underscores the significance of considering several factors and their interaction when assessing the impact of invasive species, especially considering that many habitats experience co-invasion by plants and herbivores and simultaneously undergo varous other disturbances including nutrient enrichment.

## INTRODUCTION

Human-mediated introduction of plant and animal species into biogeographic ranges where they did not occur before has been so pervasive globally that many ecosystems are now co-invaded by multiple alien plant and animal species (Dawson et al. 2017). The invasive alien species could interact with native species as well as with each other (Green et al. 2011). Invasive species often have negative interactions with native species through herbivory, parasitism, predation, and competition, which often impacts negatively on abundance and diversity of native plant and animal species and disrupt ecosystem processes (Mack et al. 2000, Wikelski et al. 2004, Parker et al. 2006, Oduor et al. 2010, Vilà et al. 2011, Ricciardi et al. 2013, Schirmel et al. 2016, Stephens et al. 2019), although there are instances where invasive species can facilitate native species diversity (Callaway et al. 2000, Oduor et al. 2018). However, far fewer studies have investigated effects of invader-invader interactions on attributes of recipient communities and ecosystem processes (O’Loughlin and Green 2017, Ricciardi et al. 2021).

Complex invader-invader interactions can lead to a wide range of outcomes for native communities and ecosystem processes (Grosholz 2005). Negative invader-invader interactions through competition and/or predation can minimize joint effects of invasive species on native biota (Ross et al. 2004). In other cases, invasive species do not have net effect on each other (Cope and Winterbourn 2004). Invasive species can also facilitate each other’s establishment, dominance, spread and ecological impacts, in accordance with the invasional meltdown hypothesis (Simberloff and Von Holle 1999, Ricciardi 2001, Best and Arcese 2009). Although empirical evidence of invaders modifying recipient ecosystems to the benefit of other aliens is accumulating, these interactions remain underexplored and underrepresented in heuristic models of invasion success (O’Loughlin and Green 2017). The complex interactions among invasive species warrant experimental research to identify how invasive species interact and their consequences for native communities and ecosystems (Johnson et al. 2009). Such studies may be particularly informative in aquatic ecosystems where experimental approaches to understanding the multi-scale effects of invasive species have historically been underrepresented compared to terrestrial ecosystems (Johnson et al. 2009, Havel et al. 2015). For most aquatic invaders, our knowledge regarding specific ecological impacts remains limited particularly at scales extending beyond the population level (Stephens et al. 2019). When ecological changes are associated with an aquatic invader, the direct and indirect mechanisms responsible are often unknown or confounded by other forms of environmental change, precluding identification of the invader’s role in observed shifts (Stephens et al. 2019). This situation may severely undermine our ability to forecast how future increases or decreases in invader abundances are likely to influence ecosystem conditions (Strayer et al. 2006).

It has been suggested that herbivory reinforces competition between plant species, which in turn diminishes the chance of coexistence among species by favoring species that are better competitors (Gurevitch et al. 2000). However, in spite of this prediction, in invasion ecology, the interactive effects of herbivory and competition on plant communities have been rarely assessed (Suwa and Louda 2012, Li et al. 2014, Zhang et al. 2018) as studies have typically assessed single biotic mechanisms (Santamaría et al. 2021). This hinders accurate estimation of the relative significance of competition and herbivory in structuring plant communities in invaded habitats, particularly in aquatic ecosystems that remain understudied (Santamaría et al. 2021).

Many freshwater ecosystems are co-invaded by aquatic macrophytes and mollusks (Tricarico et al. 2016) and at the same time experience nutrient enrichment from various sources including agricultural activities in watersheds (Havel et al. 2015). However, few studies have investigated whether effects of the multiple invaders are additive, synergistic, or antagonistic (Havel et al. 2015). Observations from field surveys that were conducted mostly in terrestrial ecosystems have found that invasive plant species generally thrive in nutrient-rich habitats (Funk and Vitousek 2007, Buckley and Catford 2016). Factorial experiments have also found that nutrient enrichment favours growth performance of invasive plant species over native plants (González et al. 2010, Liu et al. 2017). These findings generally support the suggestion that nutrient enrichment may promote plant invasion (Davis et al. 2000, Dawson et al. 2012). Theory predicts that herbivores that share evolutionary history with particular host plants will have a lower feeding preference for those plants than for plants with which they have not co-evolved (Colautti et al. 2004, Parker et al. 2006). This prediction is premised on the idea that host plants that do not share evolutionary history with herbivores have not evolved strong defences against the herbivores. Indeed, a previous synthesis of 11 studies found support for the prediction (Oduor et al. 2010). However, studies are lacking that test how nutrient enrichment and co-invasion by alien herbivores and alien plant species can interactively affect native plant communities in aquatic habitats.

Some aquatic habitats in China have been co-invaded by the apple snail *Pomacea canaliculata* and a macrophyte *Myriophyllum aquaticum* (Qiu and Kwong 2009), which are both native to South America (Hayes et al. 2008, Gillard et al. 2017), and co-occur in large areas of their native region (for details see the GBIF database; www.gbif.org). *Pomacea canaliculata* is one of the 100 malignant invasive species in the world (Lowe et al. 2000), which has become established in many tropical East Asian countries (Hayes et al. 2008). It often feeds on submerged and emergent macrophytes (Qiu and Kwong 2009) and causes economic losses and alters wetland ecosystems in its introduced region (Carlsson et al. 2004, Hayes et al. 2008). On the other hand, the macrophyte *M. aquaticum* was initially introduced as a wetland ornamental plant and sewage treatment plant (Cui et al. 2021), but spread quickly and occupied large wetlands and shallow water areas later (i.e. become invasive; Wang et al. 2016). Separate studies have shown that nutrient enrichment can enhance growth of the invasive macrophyte (Xie et al. 2010, Shen et al. 2019, Zhang et al. 2021). Studies have also shown that the invasive snail exhibits feeding preference for plants with lower defences (Qiu and Kwong 2009). Furthermore, the invasive plants often have higher resistance to generalist herbivore over native plants due to their novel allelochemicals (Schaffner et al. 2011, Qi et al. 2020). Therefore, the aquatic habitats offer a good system to test the combined effects of nutrient enrichment and co-invasion by alien herbivores and alien plant species on native plant communities.

Using a mesocosm experiment, we experimentally investigated the individual and combined effects of nutrient enrichment and co-invasion by the invasive herbivore *P. canaliculata* and the invasive plant *M. aquaticum* on biomass yield of a synthetic community of three native macrophyte species. Specifically, we tested the following predictions: (1) Nutrient enrichment enhances growth of the invasive macrophyte more than that of native macrophytes; (2) the invasive herbivore consumes more biomass of the native macrophyte than that of the invasive macrophyte because the invasive macrophyte has with a stronger defence against the herbivore, while the native macrophytes are evolutionarily naïve to the herbivore; (3) Preferential feeding by the invasive herbivore on competitor native macrophytes confers growth advantage to the invasive macrophyte.

## MATERIAL AND METHODS

### Study location and species

The study was conducted in a mesocosm of the Dongting Lake Wetland Ecosystem Observation and Research Station (29.30°N, 150.74°E) of the Chinese Academy of Sciences in Yueyang, China. The local climate is subtropical. The mesocosm comprises a fixed steel structure, with a transparent acrylic plate placed on the top. Light and temperature conditions in the mesocosm are similar to that of the ambient conditions. The roof of the mesocosm structure is covered with a waterproof nylon bag to keep out rain water. The mesocosm was subdivided into 128 individual ponds that each measured 1 m^3^.

We constructed a three-species native macrophyte community using perennial herbaceous species that commonly occur in the local wetland habitat *Vallisneria natans, Hydrilla verticillata*, and *Myriophyllum spicatum*. To simulate natural invasion of the native macrophyte community, we introduced *M. aquaticum* to the submerged native community. We procured seedlings of all the four macrophyte species from an aquatic plant-producing company (Guangzhou Beishanshui Ecological Technology Co., Ltd, Ezhou, China). The seedlings were kept in pond water until use in the experimental set up described below. On 5^th^ July 2020, we collected individuals of the apple snail from a pond around the research station. The snails were maintained in pond water with similar conditions as the experimental mesocosm for two weeks where they were fed daily with leaf tissues of the four macrophyte species that were used in the current study. Water in the pond that harbored the snails was changed daily.

### Experimental design

On 3^rd^ June 2020, we added soil into each of the 128 ponds to a depth of 15 cm. The soil was obtained from the experimental station and was sifted to remove stones and plant and animal debris. We used terrestrial topsoil instead of lake bottom silt to avoid the risk of undesired macrophytes emerging from an aquatic plant seed bank. Immediately after adding the soil, we added 150 L of groundwater to each pond. Two days later, we transplanted into the ponds similar-sized stem cuttings of all the experimental plant species except for *V. natans* that was propagated with seedlings. To facilitate transplanting, we placed a metallic framework (100 cm × 100 cm) of 36 grids (each grid measured 15 cm × 15 cm) in the center of each pond and then randomly placed 12 individual stem cuttings or seedlings of each native species in 12 grids to create a three-species native macrophyte community per pond.

Using the 128 ponds that all planted with native macrophyte community, we employed a fully crossed factorial experiment with two levels of competition from an invasive macrophyte *Myriophyllum aquaticum* (competition vs. no-competition), two levels of nutrient enrichment (enrichment vs. no-enrichment) treatment, and two levels of herbivory by an invasive snail *Pomacea canaliculata* (herbivory vs. no-herbivory) treatments. To simulate invasion of an established native macrophyte community by the invasive macrophyte *M. aquaticum*, we introduced 12 individuals of *M. aquaticum* haphazardly into half of the 128 ponds (i.e. 64 ponds; **Fig.1**) on 5^th^ June 2020. After transplanting all plants, we removed the metal frame from the ponds.

**Figure 1.**
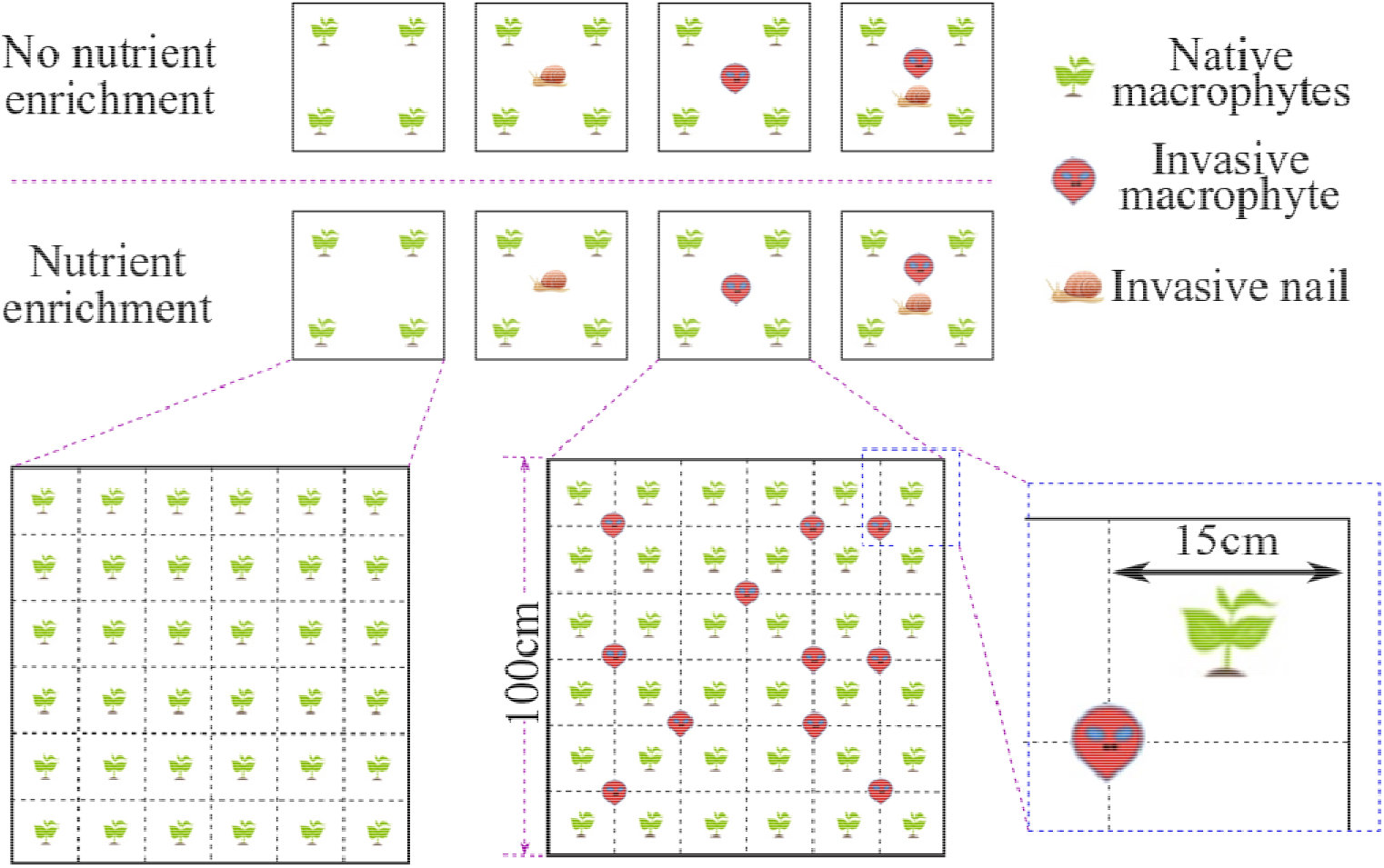
A schematic of an experimental set up to test the main and interactive effects of co-invasion by an alien snail *Pomacea canaliculata* and an alien macrophyte *Myriophyllum aquaticum* and nutrient enrichment on growth performance of a three-species community of native macrophytes (*Vallisneria natans, Hydrilla verticillata*, and *Myriophyllum spicatum*) and the invader *M. aquaticum*. Each grid represents a 1m^3^ pond; a total of 128 ponds were used in the experiment. Grids with a blue outline represent ponds with no-nutrient enrichment treatment, while grids with a red outline represent ponds with a nutrient-enrichment treatment. Six similar-sized individuals of *P. canaliculata* were introduced into each pond for a total of 64 ponds, while the other 64 ponds served as a control (no-herbivory). Nutrient-enrichment treatment was applied to 64 ponds (32 with herbivory and 32 with no-herbivory), while another 64 ponds served as a control and did not receive nutrient enrichment (32 with herbivory and 32 with no-herbivory).

On 28^th^ June 2020, we added additional 450 L of water to attain a water depth of 45 cm, and immediately thereafter imposed a nutrient enrichment treatment. We added 4.5 g of Peters Professional® water-soluble fertilizer (Total Nitrogen - 20%; Available Phosphate - 20%; Soluble Potash - 20%; Magnesium - 0.05%; Boron - 0.0125%; Copper - 0.0125%; Iron - 0.05%; Manganese - 0.025%; Molybdenum - 0.005%; Zinc - 0.025%) to a half of the ponds (i.e., 64 ponds). The fertilizer was dissolved in five liters of water and then gradually released into the pond to ensure its even distribution across the pond. As a control for nutrient enrichment, we added five liters of water to the other 64 ponds. On 13^th^ July, 2020, we introduced six similar-sized (diameter: 2.5 to 3.5 cm) individuals of the apple snail into each pond for a half the ponds. We let only the introduced snails to feed; hence, we cleared the pond of snail eggs every three days for the two-week period when herbivory treatment was imposed to prevent a new generation of snails from feeding on the macrophytes. During the first week, three dead snails from three ponds were replaced.

On 2^nd^ August, 2020, we ended the experiment when some individuals of *M. spicatum* started to flower. We removed all the apple snails and then harvested aboveground biomass of the experimental macrophytes separately for each pond. Immediately after harvest, the plants were wiped dry with absorbent paper and dried to a constant biomass in an oven at 65 °C for 72 hours. Biomass was harvested from 125 ponds as three ponds were infested by an insect that consumed all the aboveground biomass, and hence were omitted from a statistical analysis. Weight of the dry biomass was then taken to the nearest 0.1g.

To test whether the invasive snail *P. canaliculata* had a lower feeding preference for the invasive macrophytes than the native macrophytes, we set up a feeding bioassay on 7^th^ July, 2020. Ten grams of above-ground parts (including leaves and stems) of each macrophyte species was placed in a conical flask that had been filled with 1L water. Then two individuals of *P. canaliculata* of the same size (diameter: 2.5cm) were added into each flask. The flask was covered with a gauze on the top part to prevent escape of the snails and ingress of any unwanted herbivore. As a control, we set up flasks with the same amount of water and above-ground biomass of the individual macrophyte species but without the snail being introduced. Each treatment and control was replicated three times. All flasks were kept in a room at a constant temperature (27 °C) for 24h after which the experiment was stopped. Immediately at the end of the feeding bioassay, we removed all the snails and dried the plant biomass at 65 °C for 72 hours and took measurements of the individual biomass samples per flaks per herbivory treatment. The biomass records were used in the statistical tests described below.

### Statistical analysis

To test the main and interactive effects of herbivory by the invasive snail *P. canaliculata*, competition by the invasive macrophyte *M. aquaticum*, and nutrient enrichment on growth performance of the native macrophyte community, we performed a three-way ANOVA. Aboveground biomass of the native macrophyte community in each pond was specified as dependent variable. The independent variables included main and all possible three-way, and two-way interactive effects of nutrient enrichment (enrichment vs. no-enrichment), competition from *M. aquaticum* (competition vs. no-competition), and herbivory by *P. canaliculata* (herbivory vs. no-herbivory). The above-ground biomass data were log-transformed to assure normality of residuals and homogeneity of variance.

To test the main and interactive effects of herbivory by the invasive snail *P. canaliculata* and nutrient enrichment on absolute and relative growth performance of the invasive macrophyte, we conducted a two-way ANOVA using a subset of data from ponds that were invaded by *M. aquaticum*. The absolute above-ground biomass of *M. aquaticum* and the proportional above-ground biomass of *M. aquaticum* relative to the whole community above-ground biomass (i.e. an indicator of relative growth performance of *M. aquaticum*) were specified as dependent variables. The independent variables included main and two-way interactive effects of nutrient enrichment and herbivory by the invasive snail. To assure normality of residuals and homogeneity of variance, the absolute above-ground biomass and the relative above-ground biomass of the invasive macrophyte were log-transformed.

We also performed a two-way ANOVA for the feeding bioassay experiment to test the main and interactive effects of herbivory by the invasive snail and plant species identity on the biomass of above-ground parts. In the ANOVA, amount of biomass of each of the four individual macrophyte species was treated as a dependent variable, while herbivory treatment (herbivory *vs* no-herbivory) and species identity (*M. aquaticum, V. natans, H. verticillata* and *M. spicatum*) were specified as an independent variable. Then, we did a post-hoc analysis with Tukey’s Test to test whether the invasive snail had a lower preference for the invasive macrophyte than for the native macrophytes. All analyses were performed in R 4.0.3 (R Core Team 2020).

## RESULTS

Nutrient enrichment increased the above-ground biomass of the native macrophyte community (**Fig. 2; Table S1**). However, herbivory by the invasive snail significantly decreased the above-ground biomass of the native macrophyte community (**Fig. 2; Table S1**). The negative effect of herbivory by the snail was stronger when the macrophyte community was grown under nutrient-enrichment treatment than in the absence of nutrient enrichment (**Fig. 2**; significant N × H effects in **Table S1**).

**Figure 2.**
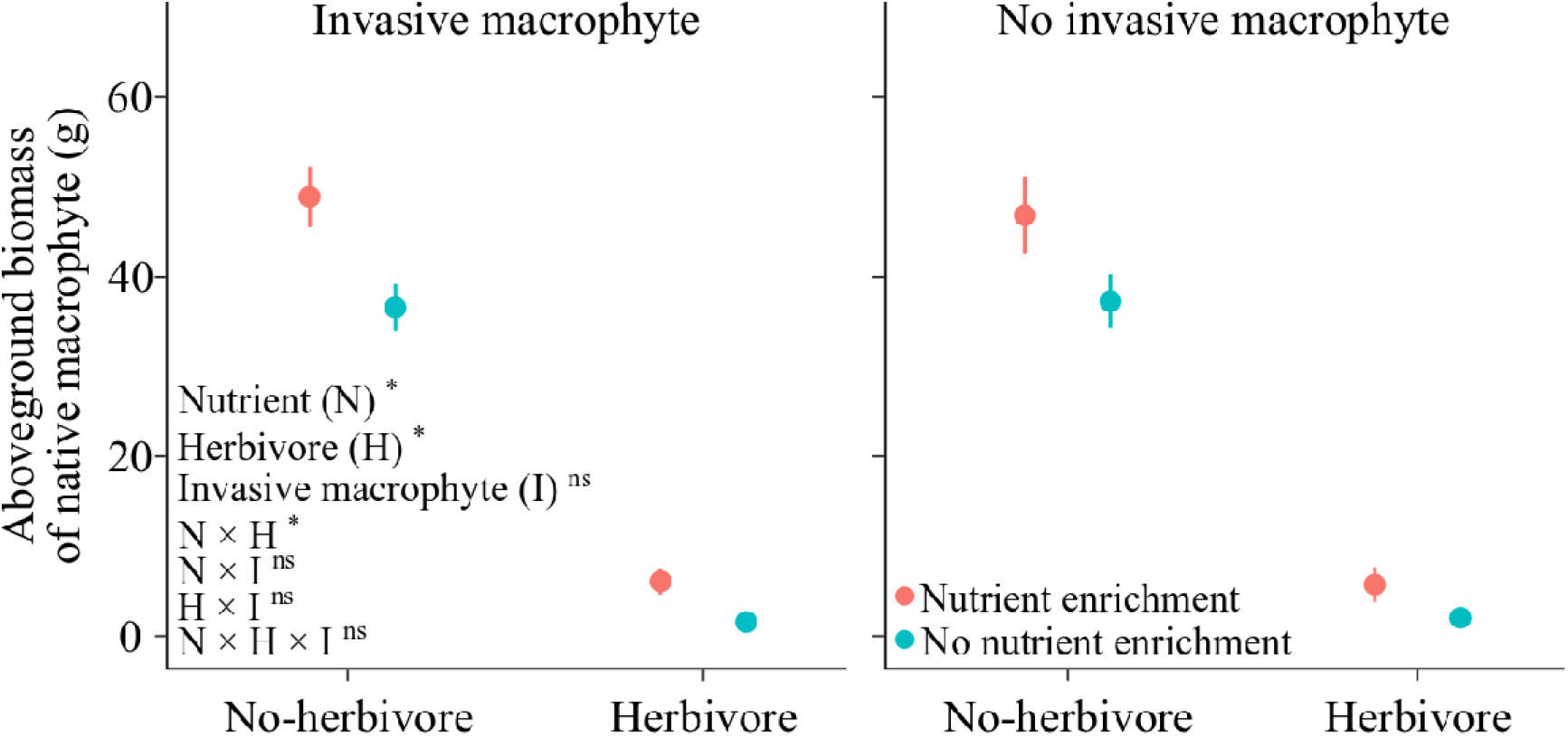
Mean (± 1 SE) above-ground biomass of a three-species native macrophyte community. The panels show the effects of invasion by an aquatic macrophyte *Myriophyllum aquaticum* under two levels of nutrient enrichment (enrichment vs. no-enrichment) and herbivory (herbivory vs. no-herbivory) by an invasive snail *Pomacea canaliculata*. The asterisks indicate level of statistical significance: * denotes *p* < 0.05, while ^ns^ denotes *p* > 0.05.

Herbivory by the invasive snail tended to decrease above-ground biomass of the invasive macrophyte (**Fig. 3a;** marginal significant effect [*p* = 0.083] in **Table S2**). However, herbivory by the invasive snail significantly increased proportional biomass of the invasive macrophyte in the invaded community (**Fig. 3b; Table S2**). Although nutrient enrichment did not affect the absolute above-ground biomass of the invasive macrophyte, nutrient enrichment decreased the proportional above-ground biomass of the invader (**Fig. 3b; Table S2**).

**Figure 3.**
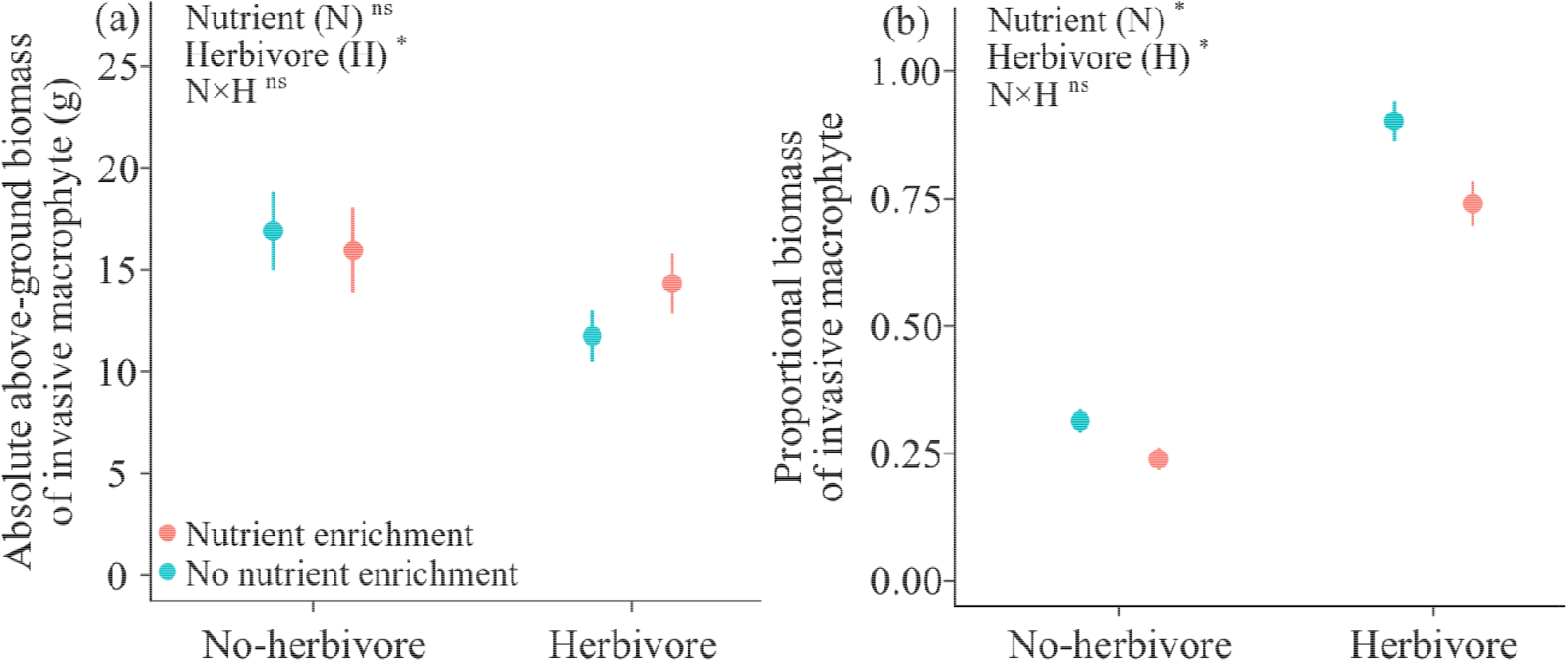
Mean (± 1 SE) absolute (**a**) and proportional (**b**) above-ground biomass of an invasive macrophyte *Myriophyllum aquaticum* when grown in a three-species native aquatic macrophyte community (*Vallisneria natans, Hydrilla verticillata*, and *Myriophyllum spicatum*). The asterisk indicate level of statistical significance: * denotes *p* < 0.05, while ^ns^ denotes *p* > 0.05.

In the feeding assay, herbivory by the invasive snail reduced significantly biomass of two of the native macrophytes *H. verticillata* and *V. natans* (**Fig. 4; Table S3**). However, biomass of a native macrophyte *M. spicatum* and the invader *M. aquaticum* was not reduced significantly by herbivory (**Fig. 4**).

**Figure 4.**
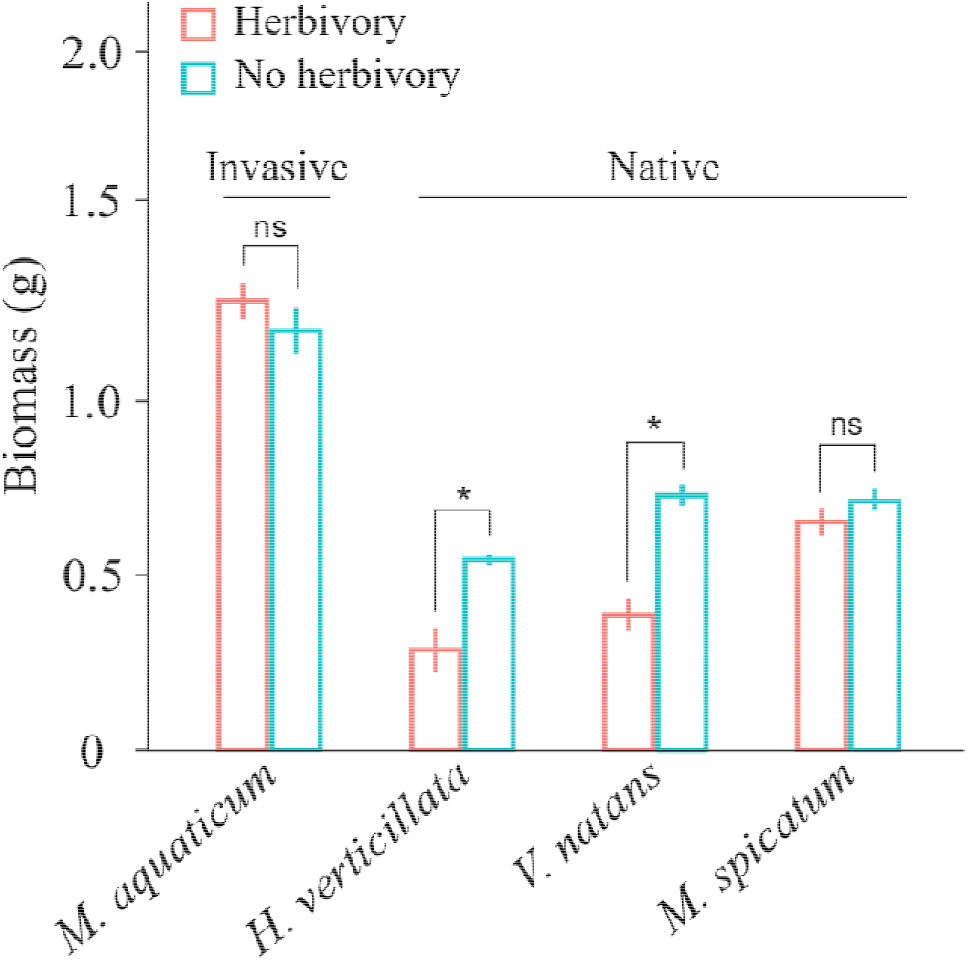
Mean (± 1 SE) biomass of three native aquatic macrophytes (*Vallisneria natans, Hydrilla verticillata*, and *Myriophyllum spicatum*) and an invasive macrophyte *Myriophyllum aquaticum* that were subjected to herbivory or not (control) by an invasive aquatic snail *Pomacea canaliculata*. Symbols above bars (*: *p* < 0.05; ^ns^: *p* > 0.05) show levels of differences between feeding and no feeding treatment based on the post-hoc analysis with Tukey’s Test.

## DISCUSSION

In line with the invasional meltdown hypothesis, we found that herbivory by the invasive snail may enhance the dominance of the invasive macrophyte in the invaded communities. On the other hand, our study does not support the prediction that nutrient enrichment may enhance invasion by the invasive macrophyte, although invasive plant species have been shown to thrive in nutrient-rich habitats in many studies (Funk and Vitousek 2007, Seabloom et al. 2015). These results generally underscore the significance of considering several factors and their interaction when assessing the impact of invasive species, especially considering that many habitats experience co-invasion by plants and herbivores (Green et al. 2011, Dawson et al. 2017) and simultaneously undergo varous other disturbances including human-induced nutrient enrichment (Bobbink et al. 2010).

Alien herbivores can facilitate invasion by alien plant species through preferential feeding on herbivory-intolerant native plants in the invaded communities (Hobbs 2001, Parker et al. 2006, Oduor et al. 2010). Indeed our results show that the invasive snail consumed more biomass of two of the three native macrophytes than that of the co-introduced invader *M. aquaticum*. As the native macrophyte species are evolutionarily naïve to the herbivore, it is likely that the native macrophytes have not evolved strong defences against the invasive snail. On the other hand, because the invasive macrophyte is native to the same biogeographic region as the invasive snail, it is likely that the invasive macrophyte has evolved strong defence against the herbivore, and hence is less palatable to the invasive snail than native macrophytes. In fact, a previous feeding assay with the invasive snail *P. canaliculata* and ten aquatic macrophyte species that included *M. aquaticum* found that the invasive snail had low survivorship and growth rate, and did not reproduce when fed on the invasive macrophyte *M. aquaticum* (Qiu and Kwong 2009). In the previous study, macrophytes with high nutritional contents and low chemical defences (i.e., low phenolic content) were more palatable to the invasive snail than macrophytes with low nutritional content and high chemical defences (Qiu and Kwong 2009). Future studies may test whether the three native macrophytes that were studied presently differ in nutritional content and chemical defences from the invasive macrophyte.

Herbivory can enhance growth performance of alien plants over native plants by inducing compensatory growth and enhanced competitive ability of alien plants (Best and Arcese 2009). Given that invasive plant species often grow faster than native species (van Kleunen et al. 2010), greater compensatory growth might be expected for the former. Plants can defend against herbivory through two strategies: resistance (i.e., plant traits that minimize damage from herbivores, e.g. defence compounds and leaf toughness) and tolerance (i.e., plant traits that enable a plant to maintain fitness after damage has occurred, e.g., increased photosynthetic and growth rates) (Strauss and Agrawal 1999, Stowe et al. 2000). Resistance and tolerance are not necessarily mutually exclusive defence strategies as plant species often deploy both. For instance, herbivory by two folivores (*Lema daturaphila* and *Epitrix parvula*) selected for plant *Datura stramonium* genotypes with intermediate resistance and high tolerance (Carmona and Fornoni 2013). Separately, the plant *Senecio jacobaea* exhibited both greater resistance to and tolerance of herbivory in the introduced range than in the native range (Stastny et al. 2005), while native populations of the invasive grass *Phragmites australis* did not display tolerance-resistance tradeoff (Croy et al. 2020). A positive association between tolerance and induced chemical defence was demonstrated in plant *Arabidopsis thaliana* (Mesa et al. 2017). Therefore, our finding that the invasive macrophyte experienced a lower herbivory by the invasive snail than native macrophytes likely because that the invasive macrophyte has high resistance.

Mutualism between invaders is posited to initiate invasional meltdown by generating reciprocal, positive population-level responses that amplify invader-specific impacts. These impacts then facilitate further invasions and accelerate the overall rate of invasion (Parker et al. 1999). Nevertheless, few studies have demonstrated positive population-level effects between invaders that amplify their impacts (Havel et al. 2015, Braga et al. 2018). Therefore, future studies may investigate population-level effects of any positive interactions between *M. aquaticum* and *P. canaliculata*.

Many empirical studies suggest that nutrient enrichment will promote alien plant invasion (Davis et al. 2000, Leishman and Thomson 2005, González et al. 2010, Dawson et al. 2012, Seabloom et al. 2015, Liu et al. 2017). However, in the present study, nutrient enrichment reduced the proportional above-ground biomass of the invasive macrophyte. It remains unclear why our results contradict those of several other studies. A plausible explanation for the current finding may be that because the current test native macrophyte species are also invasive elsewhere in the world, they have similarly high or higher growth response to nutrient enrichment than *M. aquaticum*. The native *H. verticillata* is a naturalized species in much of Asia that has invaded aquatic habitats in the United States (Zhu et al. 2017), Brazil (Sousa 2011) and South Africa (Coetzee et al. 2009). Similarly, *M. spicatum* is also listed as an invasive plant in the United States (Moody and Les 2007) and Egypt (Ali and Soltan 2006), while *H. verticillata* is highly competitive and often dominates the invasive areas and local large plant communities (Sousa et al. 2009, Hofstra et al. 2010). Although previous studies found that nutrient enrichment enhanced growth of the invader *M. aquaticum* (Xie et al. 2010, Shen et al. 2019, Zhang et al. 2021), the effect of nutrient enrichment on competitive ability of *M. aquaticum* against native macrophytes was not investigated. Therefore, it is likely that these native macrophytes have similar or higher competitive ability than *M. aquaticum* under nutrient enrichment.

In conclusion, our study suggests that nutrient enrichment in freshwater lakes may slow down invasion by the invasive macrophyte, while the inasive herbivore may enhance dominance of the invasive macrophyte in the invaded community. These results broadly support the idea that co-invasion by alien species that come from the same biogeographic region and possibly share evolutionary history can be detrimental to recipient native communities. To date, most documented cases of facilitation of one invader by another invader are from terrestrial habitats (Braga et al. 2018), and our study aims to help fill the knowledge gap of positive interactions among invasive species in aquatic habitats. Identification of interaction networks among invasive species offers opportunities beyond a single-species approach for managing invasive species in multiply invaded systems (Bull and Courchamp 2009). In the present context, dissolution of positive interactions between the invasive snail *P. canaliculata* and the invasive macrophyte *M. aquaticum* may mitigate negative impacts of the invasive macrophyte on native macrophyte communities.

## ACKNOWLEDGEMENTS

YL acknowledges funding from the Chinese Academy of Sciences (Y9B7041001). AO acknowledges funding from the CAS President’s International Fellowship Initiative (2021VBB0004).

## AUTHOR CONTRIBUTIONS

YL conceived the idea and designed the experiment. YY, FL and YX performed the experiment. YY and YL analyzed the data. YY, YL and AMO wrote the first draft of the manuscript, with further inputs from FL and YX.

## DATA ACCESSIBILITY

Should the manuscript be accepted, the data supporting the results will be archived in Dryad and the data DOI will be included at the end of the article.

## Supporting information

**Table S1.** Results of a three-way ANOVA that tested for the effects of nutrient enrichment (enrichment vs. no-enrichment), herbivory by an invasive snail *Pomacea canaliculata* (herbivory vs. no-herbivory), and presence of an invasive macrophyte *Myriophyllum aquaticum* (present vs. absent) and their interaction on above-ground biomass of a community three native aquatic macrophytes (*Vallisneria natans, Hydrilla verticillata*, and *Myriophyllum spicatum*). Significant effects (*P* < 0.05) are shown in bold font.

| Factor | Df | F | *p* |
| --- | --- | --- | --- |
| Nutrient enrichment (N) | 1 | 17.03 | **<0.001** |
| Herbivory (H) | 1 | 89.18 | **<0.001** |
| Plant invasion (I) | 1 | 0.46 | 0.139 |
| N × H | 1 | 10.34 | **0.011** |
| N × I | 1 | 1.97 | 0.419 |
| H × I | 1 | 0.37 | 0.703 |
| N × H × I | 1 | 1.53 | 0.141 |
| Residuals | Df | Mean Squares |  |
|  | 120 | 24.7 |  |

**Table S2.** Results of two-way ANOVAs that tested the effects of nutrient enrichment (enrichment vs. no-enrichment), herbivory by an invasive snail *Pomacea canaliculata* (herbivory vs. no-herbivory), and interaction between them on absolute above-ground biomass and proportional biomass of an invasive macrophyte *Myriophyllum aquaticum* when grown together with a community of three native macrophyte species (*Vallisneria natans, Hydrilla verticillata*, and *Myriophyllum spicatum*). Significant effects (*P* < 0.05) are shown in bold font.

| Factor | Absolute above-ground |  |  | Proportional above-ground biomass |  |  |
| --- | --- | --- | --- | --- | --- | --- |
|  | Df | F | *p* | Df | F | *p* |
| Nutrient enrichment (N) | 1 | 0.25 | 0.616 | 1 | 12.66 | **0.001** |
| Herbivory (H) | 1 | 3.12 | 0.083 | 1 | 212.06 | **<0.001** |
| N × H | 1 | 1.92 | 0.171 | 1 | 0.27 | 0.603 |
| Residuals | Df | Mean Squares |  | Df | Mean Squares |  |
|  | 57 | 0.090 |  | 57 | 0.205 |  |

**Table S3.** Results of a two-way ANOVA that tested for the effect of herbivory by an invasive alien snail *Pomacea canaliculata* on above-ground biomass of an invasive alien macrophyte *Myriophyllum aquaticum* and three native macrophyte species *Vallisneria natans, Hydrilla verticillata*, and *Myriophyllum spicatum*. Species identity of the four macrophytes and herbivory by the snail were treated as independent variables.

| Factors | Df | F | *p* |
| --- | --- | --- | --- |
| Species | 1 | 53.70 | **<0.001** |
| Feeding | 1 | 25.29 | **<0.001** |
| Species × Feeding | 1 | 8.18 | **0.002** |
| Residuals | Df | Mean Squares |  |
|  | 16 | 0.027 |  |

## Notes

### Competing Interest Statement

The authors have declared no competing interest.

